# Microbiota instruct IL-17A-producing innate lymphoid cells to promote skin inflammation in cutaneous leishmaniasis

**DOI:** 10.1101/2021.06.08.447514

**Authors:** Tej Pratap Singh, Augusto M. Carvalho, Elizabeth A. Grice, Phillip Scott

## Abstract

Innate lymphoid cells (ILCs) comprise a heterogeneous population of immune cells that maintain barrier function and can initiate a protective or pathological immune response upon infection. Here we show the involvement of IL-17A-producing ILCs in microbiota-driven immunopathology in cutaneous leishmaniasis. IL-17A-producing ILCs were RORγt^+^ and were enriched in *Leishmania major* infected skin, and topical colonization with *Staphylococcus epidermidis* before *L. major* infection exacerbated the skin inflammatory responses and IL-17A-producing RORγt^+^ ILC accumulation without impacting type 1 immune responses. IL-17A responses in ILCs were directed by *Batf3* dependent CD103^+^ dendritic cells, and experiments using ILC deficient *Rag1*^*-/-*^ mice established that IL-17A^+^ ILCs were sufficient in driving the inflammatory responses. As depletion of ILCs or neutralization of IL-17A diminished the microbiota mediated immunopathology. Taken together, this study indicates that the skin microbiota promotes RORγt^+^ IL-17A-producing ILCs, which augment the skin inflammation in cutaneous leishmaniasis.

## Introduction

Cutaneous leishmaniasis includes a spectrum of diseases ranging from a single ulcerative lesion to severe metastatic lesions [1]. While control of these intracellular parasites is dependent upon the production of IFN-γ by CD4^+^ T cells, the magnitude of the disease is often influenced by factors other than the parasite burden. For example, even though *L. braziliensis* patients with classical cutaneous leishmanaisis mostly control the parasites, they develop chronic lesions that can be difficult to treat [2–4]. Several studies indicate that the magnitude of disease is often due to an uncontrolled inflammatory response, which can be mediated by IL-17A and/or IL-1β [1,5–11]. Using a combination of murine models and human studies we and others have shown that the skin microbiome enhances IL-1β and IL-17A production and contributes to increased pathology in cutaneous leishmaniasis [10,12,13]. However, while the role of T cells in promoting an increased inflammatory response is well established [1], whether innate cells initiate and/or amplify a pathogenic response in leishmaniasis is unknown.

Innate lymphoid cells (ILCs) comprise a family of lymphocytes, including ILC1s, ILC2s and ILC3s, that are quickly activated by multiple soluble signals for a rapid response to infection [14,15]. ILC1s produce IFN-γ in response against pathogens, while ILC2s produce IL-5 and IL-13 in response to allergic reactions. ILC3s mainly produce IL-17 and IL-22 and have important roles in epithelial tissue repair and inflammation [14]. ILCs have been extensively studied in the gut, and recently they have been shown to play a role in the pathology in the skin [16]. For example, ILC2s contribute to atopic dermatitis, while ILC3s are present in psoriatic lesions [16–19]. Although ILC3s secrete many of the same cytokines as Th17 cells, ILC3s have distinct functional and phenotypic features and respond in an antigen independent manner [14]. Importantly, microbial products and the local cytokine environment, such as IL-23 and IL-1α and/or IL-1β produced by myeloid cells, rapidly activate ILC3s to secrete IL-17 following bacterial exposure [14,20,21]. Thus, it seemed likely that ILCs might contribute to the pathologic host-microbiota interactions in cutaneous leishmaniasis.

Recent studies suggest that alterations to the skin microbiota, particularly changes in the dominance of *Staphylococcus* and *Streptococcus* species, at the site of *Leishmania* infection are linked to disease outcome [10]. Infection-induced alterations in the skin microbiota of *L. major* infected mice are linked to disease severity and immune-mediated inflammatory responses [10,22]. Additionally, *L. major* infection of germ-free (GF) mice results in smaller lesions and reduced immunopathology relative to specific pathogen free (SPF) mice [13]. Subsequent studies found that colonization with *Staphylococcus epidermidis* promoted the generation of IL-17 producing T cells [23], which in cutaneous leishmaniasis are associated with increased disease. However, the potential role of ILCs as a source of IL-17 in cutaneous leishmaniasis has not been explored.

To determine the role of ILCs in promoting pathology we investigated if *L. major* infection increased the number of IL-17-producing ILCs in the skin following infection. While infection did not induce early IL-17 production from T cells, we found an increase in IL-17 producing RORγt^+^ ILCs following infection. Furthermore, we found that topical colonization with murine or human skin isolates of *Staphylococcus* is associated with an increase in IL-17 producing ILCs and exacerbates immunopathology in *L. major* infected mice without altering type 1 immune responses. A critical role for IL-17 producing ILCs is suggested by our finding that these ILCs are able to promote increased pathology in the absence of T cells. Taken together, these studies indicate that the skin microbiota promotes accumulation of ILC3s that exacerbate IL-17-driven immunopathology in cutaneous leishmaniasis.

## Results

### IL-17-producing RORγt^+^ILCs are enriched in skin after *L. major* infection

IL-17 has been shown to play an important role in mediating inflammation in cutaneous leishmaniasis [8,9,24] and while Th17 cells are a source of IL-17, it is also possible that IL-17 produced by ILCs present in skin could contribute to immunopathology. To assess the potential role of IL-17 from ILCs early on in *L. major* infected skin, we analyze IL-17 from skin ILCs and T cells one-week post infection **(Figure 1; Fig. S1)**. In our analysis of the cytokine IL-17 following *L. major* infection, we found no differences in IL-17 production from γδ^hi^ T cells (DETCs), γδ^low^ T cells and α β T cells compare to control mice at week one post infection **(Figure 1A and 1B** and not shown). However, the ILC3 signature cytokine IL-17 was increased in ILCs of *L. major* infected mice compared to non-infected mice **(Figure 1C and 1D)**. RORγt is expressed by ILC3s that produce IL-17 (Kobayashi et al., 2019). To determine whether these IL-17-producing ILCs in *L. major* infected skin were RORγt^+^, we examined IL-17 production from RORγt^+^ ILCs and RORγt^-^ ILCs. We found that almost all IL-17 produced by RORγt^+^ ILCs and the number of RORγt^+^ ILCs was significantly high in *L. major* infected compare to control mice **(Figure 1E and 1F)**. Consistently, we also found significant increase in the number of RORγt^+^ IL-17A^+^ ILCs in *L. major* infected compare to control mice one-week post infection **(Figure 1G)**. Moreover, we did not see induction of IFN-γ and IL-13 production from ILCs **(Figure S2A and S2B)**. We also did not see any difference in IFN-γ production from γδ^low^ T cells and α β T cells in the lesions following *L. major* infection compared to uninfected mice at week one **(Figure S3A and S3B)**. Together, these data suggest that RORγt^+^ILCs are an early source of IL-17 in the skin following acute infection by *L. major* and may contribute to the immunopathology of cutaneous leishmaniasis.

**Figure 1.**
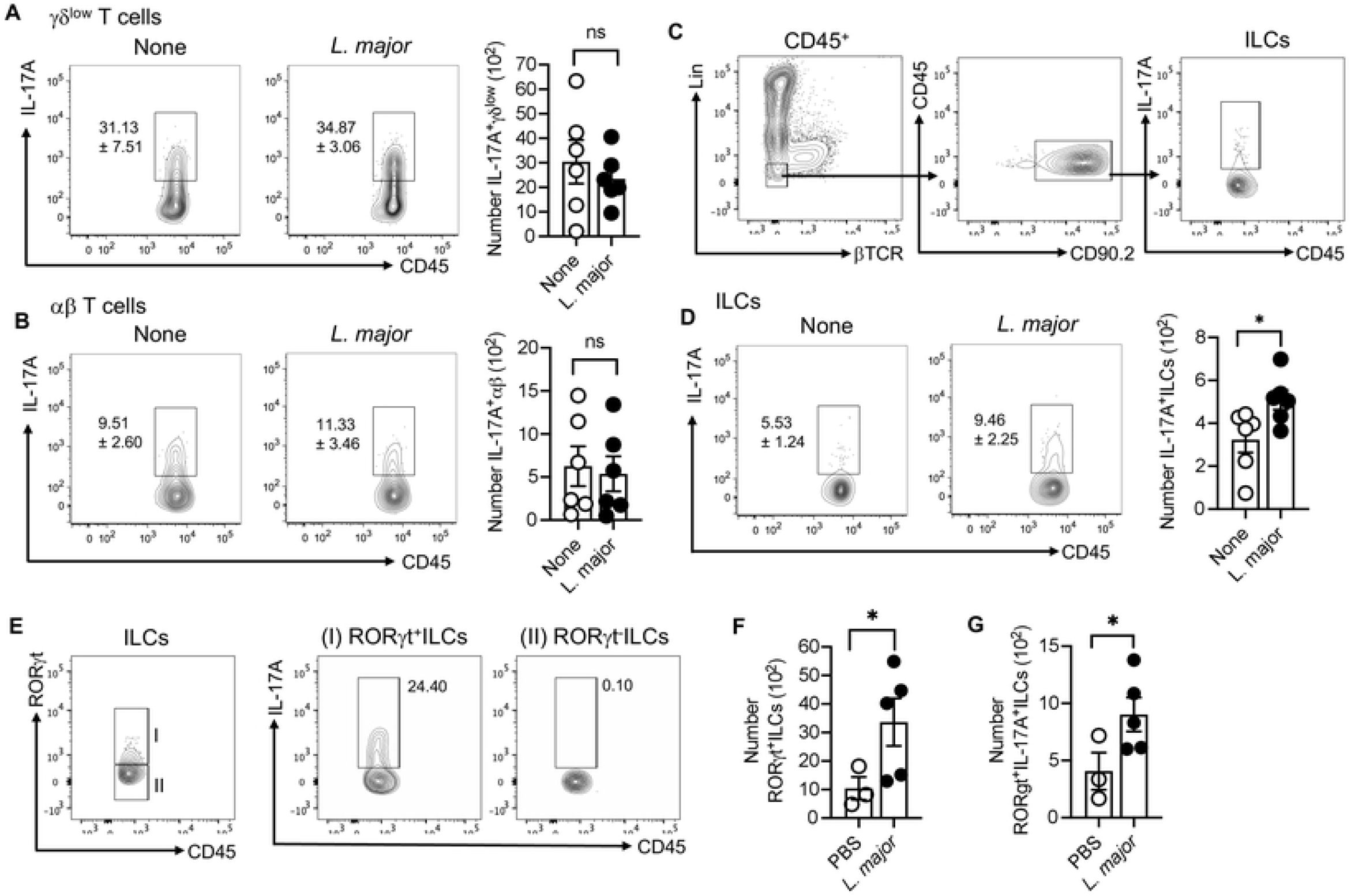
*L. major* infected skin contains IL-17A-producing RORγt^+^ILCs. **(A, B)** Percent and number of 17A+ γ δ ^low^ T cells in the skin of uninfected (None) and *L. major* infected mice at week one. **(B)** Percent and number of IL-17A+αβT cells in the skin of uninfected (None) and *L. major* infected mice at week one. **(C)** Together with Figure S1, gating strategy to identify IL-17A^+^ILCs in skin. **(D)** Percent and number of IL-17A+ILCs in the skin of uninfected (None) and *L. major* infected mice at week one. **(E)** IL-17A production from RORγt^+^ILCs and RORγt^-^ILCs in *L. major* infected mice. **(F)** Number of RORγt^+^ILCs in the skin of uninfected (None) and *L. major* infected mice at week one. **(G)** Number of RORγt^+^IL-17A^+^ILCs in the skin of uninfected (None) and *L. major* infected mice at week one. For intracellular staining cells were stimulated with PMA/Ion for 4 hours. Data are from two experiments with a total of six mice in each group (A,B,D) or from one experiment representative of two with three to five mice in each grop (E,C,F,G). Number within the flow plot show percent of IL-17A^+^ cells with SEM (A,B,D) or IL-17A^+^ cells with in the gated box (E). Error bars shows SEM. Two-tailed unpaired Student’s t-test. *p<0.05, **p<0.01, ***p<0.001.

### *S. epidermidis* enhances skin inflammation and IL-17-producing RORγt^+^ILCs in *L. major* infection

The skin microbiota contribute to the severity of human inflammatory skin diseases and our previous findings revealed a link between *Staphylococcus* and more severe disease in *L. major* infected mice [10]. To determine if skin colonization with *Staphylococcus* influences immunopathology by impacting IL-17-producing ILCs, we topically applied *S. epidermidis* to the ears and back skin of the mice once daily for five days (**Figure 2A**). As we used an m-Cherry expressing *S. epidermidis*, we quantified skin colonization by counting pink CFUs prior to infection (**Figure 2B**). One day after the last treatment with *S. epidermidis*, mice were infected in the ear with *L. major* and monitored for two weeks. We found that *S. epidermidis* colonization before *L. major* infection significantly increased the inflammatory response as compared to *L. major* infected mice alone (**Figure 2C, 2D and 2E**). In contrast, *S. epidermidis* association without *L. major* infection did not elicit an inflammatory response **(Figure 2C, 2D and 2E)**. Moreover, flow cytometry analysis revealed that *S. epidermidis* colonization before *L. major* infection increased the IL-17^+^ILCs around two-fold compared to *L. major* alone (**Figure 2F and 2I**). Moreover, RORγt^+^ ILCs were significantly increased in *S. epidermidis* colonized and *L. major* infected mice compared to *L. major* infected mice alone **(Figure 2G and 2J)**. Consistently, in analyzing IL-17 production from RORγt^+^ ILCs and RORγt^-^ ILCs, we found that almost all IL-17 was produced by RORγt^+^ ILCs in *S. epidermidis* colonized and *L. major* infected mice **(Figure 2H)** and the number of RORγt^+^ IL-17A^+^ ILCs were increased in these mice compare to *L. major* alone **(Figure 2K)**. Next, we extended our findings using an isolate of *Staphylococcus xylosus* that was obtained from *L. major* infected mice that is also commonly found on uninfected murine skin and wounds [25,26]. Consistently, *S. xylosus* colonization before *L. major* infection led to an increased inflammatory response as determined by skin thickness and pathology score compared to *L. major* infected mice without colonization **(Figure 2K and 2L)**. In addition, *S. xylosus* colonization also increased the number of IL-17^+^ILCs compared to uncolonized *L. major* infected mice **(Figure 2N and 2O)**. These data suggest that murine and human skin commensals exacerbate IL-17^+^ILC -driven inflammatory responses in the skin during cutaneous leishmaniasis.

**Figure 2.**
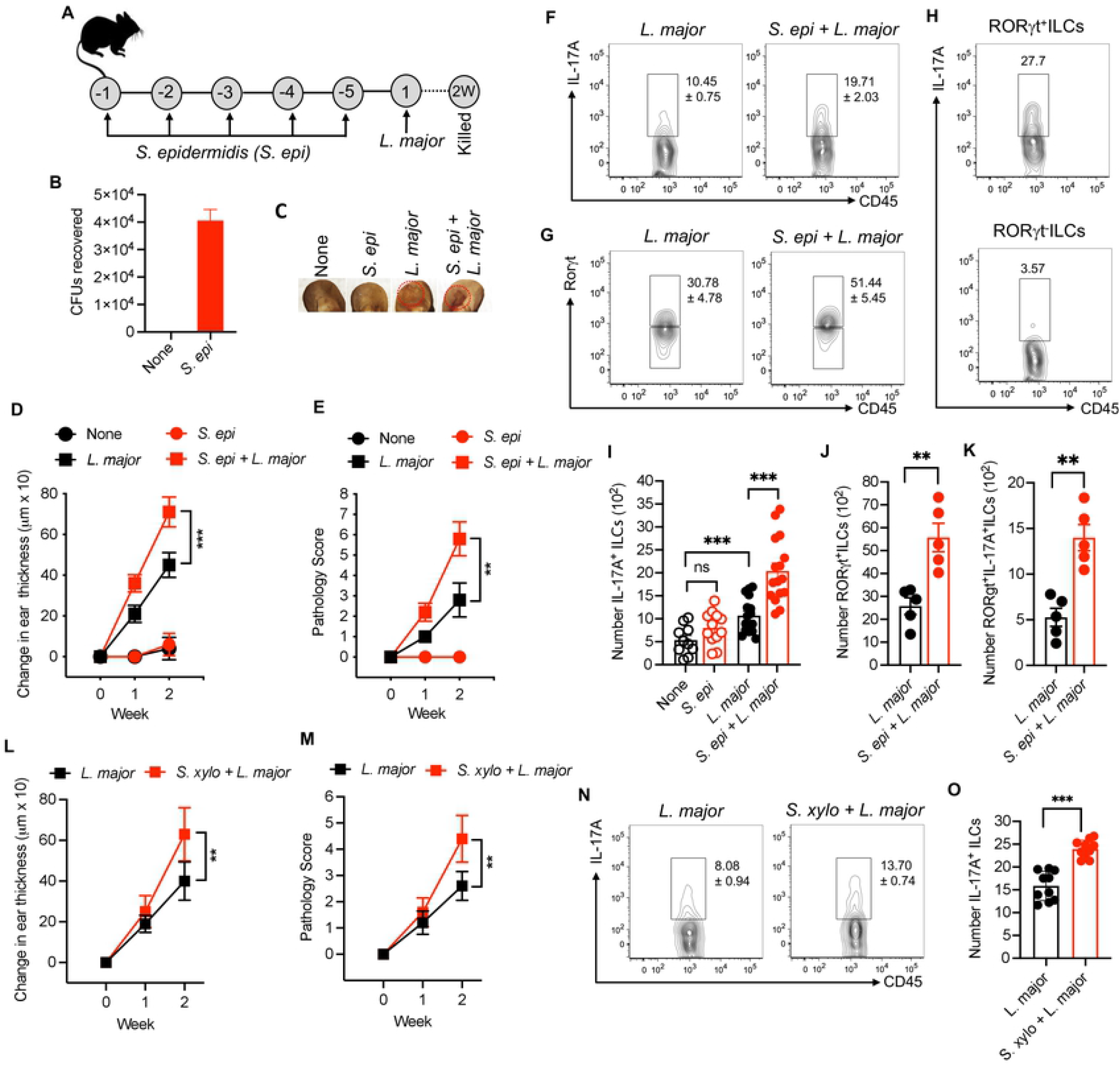
*S. epidermidis and S. xylosus* colonization before *L. major* infection increase the inflammatory responses and IL-17A^+^ILCs. **(A)** Schematic representation of *S. epidermidis* (*S. epi*) and *L. major* treatment protocol in B6WT mice. **(B)** Recovered colony forming unit (CFU) in the ear of uncolonized (None) and *S. epi* colonized mice before injecting *L. major* in the skin. **(C)** Representative ear of unassociated (None), *S. epi* associated, *L. major* infected and *S. epi* associated plus *L. major* infected mice. **(D, E)** Ear thickness measurement and pathology score in mice associated with *S. epi* or unassociated prior to *L. major* infection at week one and two. **(F)** Percent of IL-17A+ILCs in the skin of different treatment groups at week two. **(G)** Percent of RORγt^+^ILCs at week two in the skin of different treatment groups at week two. **(H)** IL-17A production from RORγt^+^ILCs and RORγt^-^ILCs in *S. epi* colonized and *L. major* infected mice. **(I)** Number of IL-17A+ILCs in the skin of different treatment groups at week two. **(J)** Percent of RORγt^+^ILCs at week two in the skin different treatment groups at week two. **(K)** Percent of RORγt^+^IL-17A^+^ILCs at week two in the skin different treatment groups at week two. **(L**,**M)** Ear thickness measurement and pathology score in mice associated with *S. xylosus (S. xylo)* or unassociated prior to *L. major* infection at week one and two. **(N)** Percent of IL-17A+ILCs in the skin of different treatment groups at week two. **(O)** Number of IL-17A+ILCs in the skin of different treatment groups at week two. Number with in the flow plot show percent of IL-17A^+^ cells with SEM (F,G,N) or IL-17A^+^ cells with in the gated box (H). Data are from three experiments with a total of 10 to 16 mice in each group (I,O) or from one experiment representative of two with five mice in each group (B,D,E,H,K,L,M). Error bars shows SEM (B,I,J,K,O) or SD (D,E,L,M). Two-tailed unpaired Student’s t-test or one-way ANOVA with Tukey’s multiple comparision analysis (I). ns, not significant; *p<0.05, **p<0.01, ***p<0.001.

### *S. epidermidis* does not impact type 1 immune responses against *L. major*

To assess if commensal bacterial colonization of the skin alters type 1 immunity in our studies, we colonized the mice with *S. epidermidis* for 5 days as described above and then infected the mice with *L. major*. At two weeks post-infection lesions were analyzed for immune responses and parasite burden. There was no difference in IFN-γ production from α β T cells in *S. epidermidis* colonized and *L. major* infeted mice compare to *L. major* infected mice alone (**Figure 3A and 3B**), and the numbers of *L. major* parasites were similar in both groups. Next, we similarly evaluated the effect of *S. xylosus* on IFN-γ production from α β T and parasite load; mice colonized with *S. xylosus* before *L. major* infection exhibited a slight decrease in IFN-γ production from α β T cells but did not show any difference in the parasite burden **(Figure 3D, 3E and 3F)**. Overall, these results suggest that the skin commensals that we used in this study does not dramatically alter the type 1 immunity against *L. major* during the acute phase.

**Figure 3.**
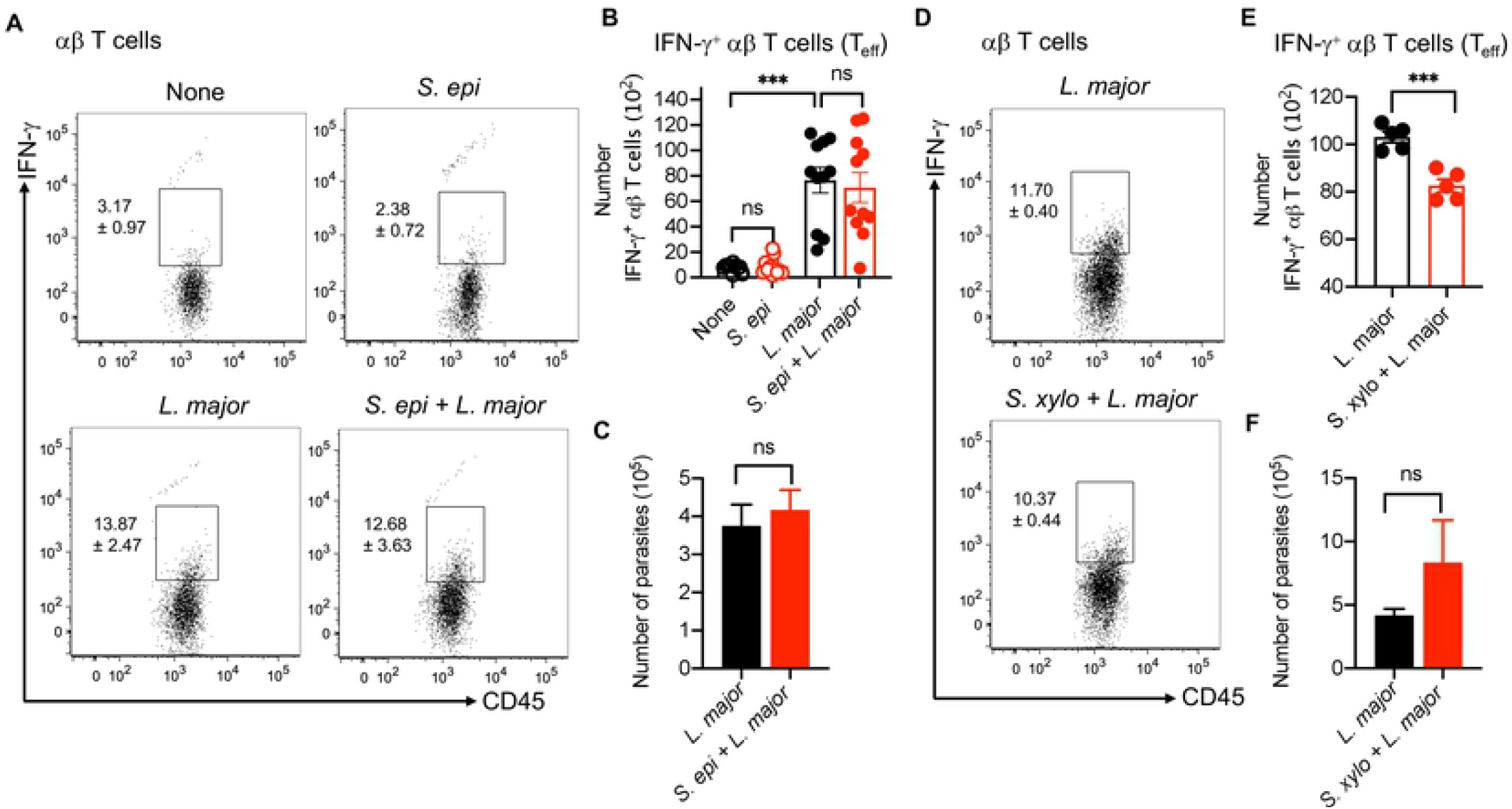
*S. epidermidis and S. xylosus* colonization does not impact the type 1 immune response. **(A)** Percent of IFN-γ-producing αβT cells in the skin of different treatment groups at week two. **(B)** Number of IFN-γ-producing αβT cells in the skin of different treatment groups at week two. **(C)** *L. major* parasite load in different treatment groups at week two. **(D)** Percent of IFN-γ-producing αβT cells in the skin of different treatment groups at week two. **(E)** Number of IFN-γ-producing αβT cells in the skin of different treatment groups at week two. **(F)** *L. major* parasite load in different treatment groups at week two. Number with in the flow plot show percent of IFN-γ^+^ cells with SEM (A,D) Data are from three experiments with a total of 10 to 12 mice in each group or from one experiment representative of two with five mice in each grop (C,D,E,F). Error bars shows SEM. Two-tailed unpaired Student’s t-test or one-way ANOVA with Tukey’s multiple comparision analysis (B). ns, not significant; ***p<0.001.

### *S. epidermidis* dependent IL-17-producing ILCs and inflammation require CD103^+^ dendritic cells

Microbes and their products are potent inducers of innate and adaptive immune cell responses driven by dendritic cells (DCs). The skin contains different subsets of DCs that drive unique immune responses, and previous reports indicate that CD103^+^ DCs in skin are the primary sensor of commensals that regulate formation of commensal-specific T-cell responses [23]. CD103^+^ DCs are classical CD11c^+^CD11b^-^ DCs that depend on the transcription factor *Batf3* for their development in the skin [23]. Therefore, we employed *Batf3*^*-/-*^ mice to explore whether or not CD103^+^ DCs are required for IL-17-producing ILC responses and enhanced immunopathology due to *S. epidermidis* colonization during *L. major* infection (**Figure 4A**). As expected, *L. major* infected *Batf3*^*-/-*^ mice, with or without colonization, lack CD103^+^ DCs in the skin (**Figure 4D and 4E**). In contrast, topical association with *S. epidermidis* increased the number of CD103^+^ DCs in skin of WT mice (**Figure 4D,E**). Although the lack of CD103^+^ DCs did not affect the lesion size in the *Batf3*^*-/-*^ mice compared to wild-type mice infected with *L. major* alone (**Figure 4B and 4C)**, the absence of CD103^+^ DCs completely abrogated the *S. epidermidis* mediated effect on immunopathology during *L. major* infection, as assessed by ear thickness and pathology score (**Figure 4B and 4C)**.

**Figure 4.**
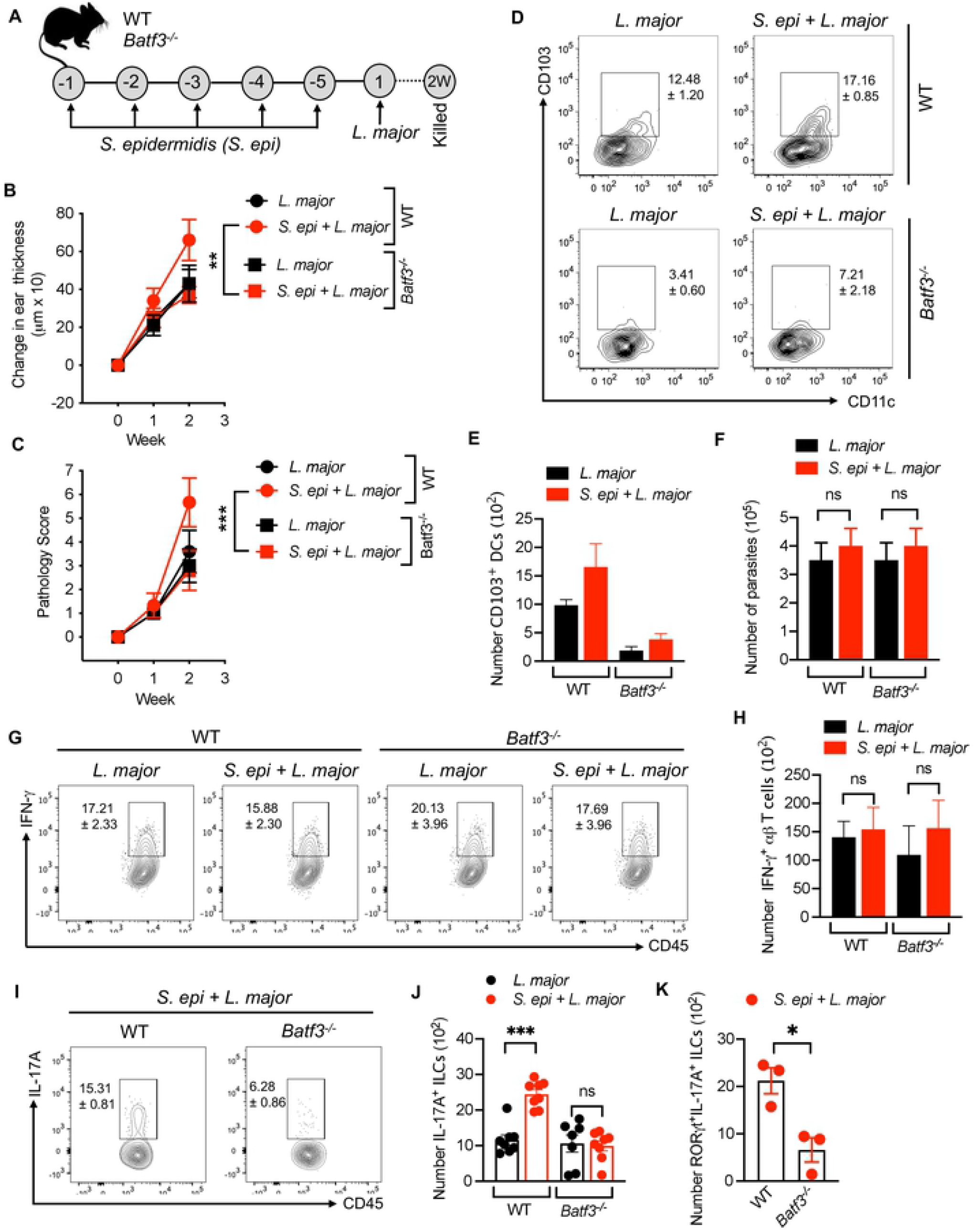
*S. epidermidis* dependent IL-17A^+^ILCs and inflammatory responses require CD103^+^ DCs. **(A)** Schematic representation of *S. epi* and *L. major* treatment protocol in WT and *Batf3*^*-/-*^ mice. **(B**,**C)** Ear thickness measurement and pathology score in WT and *Batf3*^*-*/-^ mice associated with *S. epi* or unassociated prior to *L. major* infection at week one and two. **(D**,**E)** Percent and number of CD103^+^ DCs at week two in WT and *Batf3*^*-*/-^ mice associated with *S. epi* or unassociated prior to *L. major* infection at week two. **(F)** Parasite load in WT and *Batf3*^*-*/-^ mice associated with *S. epi* or unassociated prior to *L. major* infection. **(G**,**H)** Percent and number of IFN-γ^+^αβT cells in WT and *Batf3*^*-*/-^ mice associated with *S. epi* or unassociated prior to *L. major* infection at week two. **(I**,**J)** Percent and number of IL-17A-producing ILCs in WT and *Batf3*^*-*/-^ mice associated with *S. epi* or unassociated prior to *L. major* infection. Number within the flow plot show percent of CD103^+^CD11c^+^ (D), IFN-γ^+^ (G) and IL-17A^+^ (I) cells with SEM. Data are from three experiments with a total of six to eight mice in each group (D,E,I,J) or from one experiment representative of two with three to five mice in each grop (B,C,F,K). Error bars shows SEM (E,F,H,J,K) or SD (B,C). Two-tailed unpaired Student’s t-test or one-way. ns, not significant; *p<0.05, **p<0.01, ***p<0.001.

To determine if the CD103^+^ DC-dependent effects of *S. epidermidis* colonization on *L. major* induced pathology were due to the lack of a type 1 immune response, we next evaluated the parasite burden and the number of adaptive immune cells in the lesion. CD103^+^ DCs have been shown to be critical for the production of IL-12 in leishmaniasis, and *Batf3*^*-/-*^ mice develop an uncontrolled infection over time [27]. However, similar to what has been previously reported [28], at this early time point parasite number and the number and percent of IFNγ ^+^ α β T cells in the skin of WT and *Batf3*^*-/-*^ mice were similar **(Figure 4F, 4G and 4H)** indicating that the lack of CD103^+^ DC did not impact early type 1 immune responses or control of *L. major* parasites. Furthmore, we analyzed IL-17-producing ILCs in WT and *Batf3*^*-/-*^ mice infected with *L. major* alone or in conjunction with *S. epidermidis* colonization. While there was no difference in the number of IL-17^+^ILCs in WT and *Batf3*^*-/-*^ mice infected with *L. major* alone, the number of IL-17^+^ILCs was significantly reduced in *L. major* infected *Batf3*^*-/-*^ mice colonized with *S. epidermidis* when compared to similarly treated WT mice **(Figure 4I and 4J)**. Similarly, the number of RORγt^+^ IL-17A^+^ ILCs were also decreased in *L. major* infected *Batf3*^*-/-*^ mice colonized with *S. epidermidis* compared to WT mice (**(Figure 4K**). These data suggest that *S. epidermidis* mediated IL-17-producing ILCs responses are dependent on skin CD103^+^DCs and support the idea that microbes stimulate CD103^+^ DCs to drive the induction and/or maintenance of IL-17-producing ILCs that contribute to immunopathology in cutaneous leishmaniasis.

### IL-17^+^ ILCs are sufficient in mediating *S. epidermidis* dependent inflammation

To determine if IL-17 from ILCs can drive the inflammatory responses in the absence of T cells, we colonized *Rag1*^*-/-*^ mice with *S. epidermidis* and then infected them with *L. major* or infected uncolonized mice with *L. major* **(Fig. 5A)**. Colonization with *S. epidermidis* increased the ear thickness and pathology score of *Rag1*^*-/-*^ mice compared to *L. major* infected mice alone **(Figure 5B and 5C)**. In analyzing the IL-17^+^ILCs in different treatment groups, we found that *S. epidermidis* colonization significantly increased the IL-17^+^ILCs in *L. major* infected *Rag1*^*-/-*^ mice compared to uncolonized mice **(Figure 5D and 5E)**. Of note, analysis of IFN-γ and IL-5 production in treatment groups revealed no difference in IFN-γ and reduced IL-5 from ILCs in *S. epidermidis* associated and *L. major* infected mice compared to uncolonized mice **(Figure S4A and S4B)**.

**Fig. 5.**
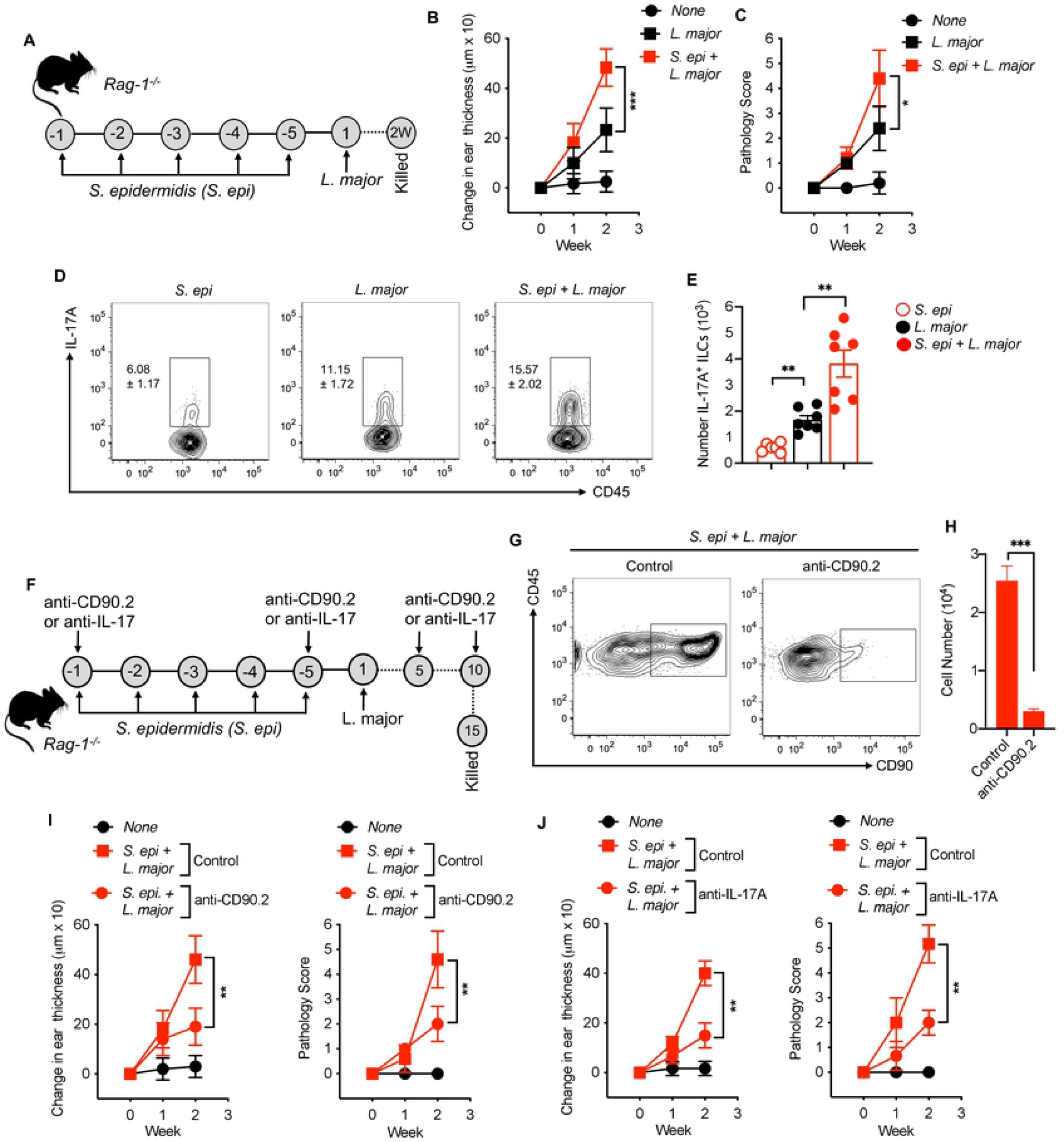
IL-17A^+^ILCs are sufficient to drive *S. epidermidis* mediated inflammation. **(A)** Schematic representation of *S. epi* and *L. major* treatment protocol in *Rag1*^*-/-*^ mice. **(B, C)** Ear thickness measurement and pathology score in *Rag1*^*-/-*^ mice associated with *S. epi* or unassociated prior to *L. major* infection at week one and two. **(D**,**E)** Percent and number of IL-17A-producing ILCs at week two in *Rag1*^*-/-*^ mice associated with *S. epi* or unassociated prior to *L. major* infection. **(F)** Schematic representation of ILC depletion or IL-17 neutralization treatment protocol in *Rag1*^*-/-*^ mice. **(G)** Representative flowcytometry plot of ILC depletion as identified as CD45^+^CD90^+^ cells in *Rag1*^*-/-*^ mice. **(H)** Number of CD90^+^ cells in control and anti-CD90.2 treated mice. **(I)** Ear thickness measurement (left) and pathology score (right) in *Rag1*^*-/-*^ mice with and without (Control) ILC depletion by anti-CD90.2 antibody associated with *S. epi* prior to *L. major* infection. **(J)** Ear thickness measurement (left) and pathology score (right) in *Rag1*^*-/-*^ mice with and without (Control) IL-17A neutralization by anti-IL-17A antibody associated with *S. epi* prior to *L. major* infection. Number with in the flow plot show percent of IL-17A^+^ cells with SEM (D) Data are from two experiments with a total of six to seven mice in each group. Error bars shows SEM (E,H) or SD (B,C,F,G). Two-tailed unpaired Student’s t-test. *p<0.05, **p<0.01, ***p<0.001.

To directly demonstrate that ILCs were promoting the increased immunopathology following *L. major* infection in *Rag1*^*-/-*^ mice, we depleted ILCs by injecting anti-CD90.2 antibody into *Rag1*^*-/-*^ mice just before the first *S. epidermidis* colonization and then at days 5 and 10 after *L. major* infection **(Figure 5F)**. Depletion of CD90.2 cells in *Rag1*^*-/-*^ mice was confirmed by flow cytometry **(Figure 5G and FH)**. Depleting CD90.2^+^ ILCs significantly reduced the inflammatory responses as assessed by ear thickness and pathology score two weeks post-infection in *S. epidermidis* colonized and *L. major* infected *Rag1*^*-/-*^ mice compared to *S. epidermidis* colonized and *L. major* infected *Rag1*^*-/-*^ mice that did not receive the anti-CD90.2 antibody (‘control’) **(Figure 5I)**. To test that IL-17 production by ILCs was responsible for mediating the pathology in *Rag1*^*-/-*^ mice, we treated mice with anti-IL-17A mAb. Notably, blockade of IL-17A in *Rag1*^*-/-*^ mice colonized with *S. epidermidis* before *L. major* infection significantly reduced the inflammation (**Figure 5J)**, suggesting that IL-17 production from ILCs is sufficient to drive inflammation in acute phase of cutaneous leishmaniasis. Together, these results support our notion that IL-17-producing RORgt^+^ILCs are critical mediators of microbiota-driven inflammatory responses in cutaneous leishmaniasis.

## Discussion

Cutaneous leishmaniasis exhibits a wide spectrum of clinical presentations, and understanding the mechanisms driving these diverse manifestations is critical for the development of new therapies. In some cases, the lack of an effective immune response leads to uncontrolled parasite replication leading to severe disease [1]. However, in other situations effective immunity develops, but an exaggerated inflammatory response sustains disease [1]. Here we report one pathway that leading to increased disease without an increase in the parasite burden is mediated by bacterial colonization and the subsequent expansion of a pathologic IL-17-producing RORγt^+^ILC population in the skin.

It appears that several pathways can lead to pathologic inflammatory responses in cutaneous leishmaniasis, including extensive cell lysis leading to inflammasome activation and IL-1 production [7,29–31], infection with more virulent strains of the parasite [6] and the lack of regulatory cytokines such as IL-10 leading to an IL-1 and IL-17 mediated pathology [32]. Additionally, alterations in the skin microbiome influence disease. We found that during *L. major* infection of mice there is a dramatic decrease in bacterial diversity in the skin that coincides with increases in *Staphylococcus* relative abundance. In co-housing experiments we demonstrated that transfer of this lesion-associated microbiota promotes increased severity of leishmanial lesions [10]. Correspondingly, germ-free mice develop smaller lesions than conventional mice, and colonization of mice with *Staphylococcus* leads to increased disease [13]. Thus, while alterations to the microbiota can be a consequence of inflammation and tissue damage, these studies demonstrate the potential for the skin microbiota to promote inflammation. Alterations to the skin microbiome occur in both murine and human cutaneous leishmaniasis [10,33], and our studies indicate how those alterations impact IL-17-producing RORγt^+^ILCs and their contributions to immunopathology in cutaneous leishmaniasis.

ILCs are a heterogeneous group of innate immune cells: ILC1s secrete IFN-γ, ILC2s secrete IL-5 and IL-13, and ILC3s secrete IL-17 and IL-22 [34]. As the role of ILCs has not been investigated in leishmaniasis, we first asked whether the acute infection altered the ILCs in the skin. Since *L. major* infection in mice that resolve the disease is associated with a CD4 Th1 response, we predicted that there might be an increase in IFN-γ from ILCs early after infection. Consistent with this prediction was our previous finding that NK cells in *L. major* infected resistant mice contribute to an early Th1 response [35]. However, we found no significant changes or induction of IFN-γ or IL-13 production from ILCs after infection, suggesting that these cytokine secreting ILCs probably are not major players in the early immune response to *L. major* in B6 mice. Whether they might play a role in other resistant strains is unknown. In contrast, we observed, an increase in ILCs producing IL-17 that was driven by *L. major* infections and further enhanced by bacteria colonization of the skin, leading to increased disease. IL-17-producing ILCs (ILC3s) are present at barrier surfaces in close contact with the microbiota, which can have a profound effect on tissue inflammation and homeostasis [14,15]. Further, it is known that the gut and skin commensal microbiota play an important role in regulating IL-17 responses in ILC3 [16,21], which is consistent with our findings. Such results indicate that the endogenous skin microbiota will likely impact responses to both infection and vaccination and that therapeutic modulation of the commensal microbiota in skin could potentially be harnessed to increase vaccine efficacy [36,37].

IL-17A has been implicated in the immunopathology of several experimental models of cutaneous leishmaniasis. Notably, *S. aureus* infection together with *L. major* exacerbates the IL-17A dependent pathology [12], and cytoplasmic virus within a strain of *L. guyanensis* induces IL-17A production to mediate diseases severity [38]. We extend the current understanding of immunopathology in cutaneous leishmaniasis by demonstrating that commensal microbiota induced IL-17A-producing RORγt^+^ ILCs that drive early pathology in *L. major* infected mice. In support of a pathogenic role for these cells in the skin, IL-17A producing RORγ t^+^ ILCs have been implicated in models of psoriasis [15,17,39]. Furthermore, the increased numbers of IL-17-producing ILC3s in lesions of psoriatic patients were positively correlated with disease severity, and adoptive transfer of ILC3s that can produce IL-17 in a human xenotransplant mouse model was sufficient to induce psoriasis [40]. By using *Rag1*^*-/-*^ mice which lack T and B cells, we directly demonstrated the importance of IL-17^+^ILCs and commensal microbiota in mediating the skin pathology in *L. major* infection, as IL-17A neutralization or CD90.2 depletion in *Rag1*^*-/-*^ mice reduced the pathology in *L. major* infected mice associated with *S. epidermidis*. Thus, our data suggest that the role of IL-17-producing RORγt^+^ ILCs in inducing pathology is not limited to inflammatory skin conditions such as psoriasis but contributes to other diseases such as cutaneous leishmaniasis.

We found that *Staphylococcus* colonization increased lesion size and the degree of pathology in mice infected with *L. major*, while not influencing the parasite load. We observed this result both with *S. epidermidis* and a strain of *S. xylosis* that we isolated from mice infected with *L. major* [10]. These results are similar to our previous findings where co-housed mice had more severe disease, but no change in the parasite burden [10]. Since the parasite burden does not change, it is not surprising that there was no alteration in the IFN-γ response. However, it is clear that in some situations the microbiota can influence IFN-γ responses. In one study the total lack of microbiota in germ-free C57BL/6 mice infected with *L. major* led to low levels of IFN-γ, suggesting that some threshold of microbiota may be required for optimal IFN-γ responses [13]. However, contradictory results were obtained in another study, where the parasite burden was higher in germ-free mice, while the levels of IFN-γ were similar between SPF and germ-free mice infected with *L. major* [41]. These differences may be due to differences in the genetic background of the germ-free mice, the route of infection, or the strain or species of leishmania studied [13,22,41].

The pathway leading to expansion of IL-17^+^ RORγt^+^ ILCs following bacterial association and *L. major* infection may be similar to that driving T cell production of IL-17. We found that *Batf3* dependent CD103^+^ DCs were required for IL-17-producing ILCs responses. Contrasting reports indicate that, relative to WT mice, *L. major* infection of *Batf3*^*-/-*^ mice develop either exacerbated [27] or similar pathology [28]. In our study, we did not observe differences in IFN-γ production from α β T cells or parasite load in *Batf3*^*-/-*^ mice compared to wild-type mice after infection. However, we did find that colonization with *S. epidermidis* in *Batf3*^*-/-*^ mice failed to induce IL-17 from RORγt^+^ ILCs or exacerbate the pathology in *L. major* infected mice. These results are consistent with reports showing a role for CD103^+^ DCs in priming innate immune cells for IL-17 production. For example, the expansion of CD8^+^ T cells producing IL-17 following colonization of mice with a *S. epidermidis* strain was dependent on CD103^+^ DCs [23]. Similarly, the bacterial component flagellin induces IL-23 production by intestinal CD103^+^CD11b^+^ DCs to activate RORγt^+^ ILC3s [42], and microbiota-activated CD103^+^ DCs drive γδT17 proliferation and activation [43]. Thus, our data show that there is a requirement for CD103^+^ DCs, but it does not exclude a contributing role for other DCs in the skin [23].

Together our data support the notion that microbiota-driven IL-17^+^RORγt^+^ILC activation can promote increased immunopathology, thus further demonstrating the role that IL-17 plays in cutaneous leishmaniasis. The importance of IL-17 in promoting disease may not be confined to murine models, as patients with classical cutaneous leishmaniasis and mucosal leishmaniasis express IL-17 in lesions [32,44]. Similarly, leishmania infection induces alterations to the skin microbiome in patients as well as mice [10,33]. Accordingly, we found *L. braziliensis* patients often had a dominant *Staphylococcus* dysbiosis, which could be a driver of increased IL-17 [10]. Taken together, these results provide a rationale for therapeutic targeting of commensals, innate lymphocytes, and IL-17 for the treatment of cutaneous leishmaniasis.

## Methods

### Mice

Male wild-type C57BL/6 mice were purchased from Charles River Laboratories (Durham, NC). Rag1^−/−^ (B6.129S7-RAG1^tm1Mom^/J) mice were purchased from The Jackson Laboratory and bred in our facility. *Batf3*^*-/-*^ (B6.129P2(C)-Batf3^tm1Kmm^/J) mice were the gift from Drs. C. Hunter and D. Herbert (University of Pennsylvania, PA.) All mice were maintained in specific pathogen-free facilities at the University of Pennsylvania. Mice were randomly assigned to experimental groups and were 6–8 wk old at the start of the experiment and were age-matched within each experiment. All procedures involving mice were performed in accordance with the guidelines of the University of Pennsylvania Institutional Animal Care and Use Committee (IACUC).

### *L. major* culture and infection

*L. major* (WHO/MHOM/IL/80/Friedlin) parasites were grown in Schneider’s Drosophila medium (GIBCO BRL, Grand Island, NY, USA) supplemented with 20% heat-inactivated fetal bovine serum (FBS) (Invitrogen USA), and 2 mML-glutamine for 4-5 days. The infectious metacyclic promastigotes of *L. major* were isolated by Ficoll (Sigma) density gradient centrifugation [45]. Mice were infected intradermally in the ear with 1×10^5^ *L. major* parasites.

### Lesion measurement and pathology score

The development of lesions was monitored weekly by measuring the ear thickness with a digital caliper (Fisher Scientific). Ear swelling/thickness was determined for individual mice by subtracting the ear thickness before treatment from that after treatment at the different time points. Inflammation and pathology were assessed by using the following inflammatory features: swelling/redness, deformation, ulceration, and loss of tissue. Based on the macroscopic appearance of the skin each feature was scored as no symptom (0), mild (1), moderate (2) and severe (3) of individual mice. The scores were summed, resulting in a maximal score of 12. Parasite burden in lesion tissues was assessed using a limiting dilution assay as previously described [10].

### *S. epidermidis* and *S. xylosus* colonization

*Staphylococcus epidermidis* strain Tu3298 expressing a fluorescent protein mCherry was a gift of Dr. Tiffany Scharschmidt (UCSF) [46] and *Staphylococcus xylosus* an isolate that was cultured from the ears of *L. major* infected mice [10] were used in the experiments. For topical association, the bacteria were cultured for 24 hours in a shaking incubator, washed and re-suspended in PBS. 10^8^-10^9^ CFUs of bacteria were applied to the back and ears of the mouse using sterile cotton swabs, every day for a total of 5 days before injecting the *L. major*. For CFU quantification with *S. epidermidis*, digested ears were plated on soy agar plate and incubated overnight at 37^0^C. The expression of pink colonies allowed us to directly visualize the bacteria.

### Depletion of ILCs and IL-17A neutralization

To deplete ILCs, Rag1^-/-^ mice were injected with 100μg anti-CD90.2 antibody (30H12; BioXCell). To study the effect of IL-17A in mediating the immunopathology, Rag1^-/-^ mice were injected with 10 mg/kg anti-IL-17A antibody (17F3; BioXCell). In both cases mice were injected with the anti-antibody one day before the start of the *S. epi* colonization at day -1 and before the *L. major* injection at day -5 and then every 5 days at day 5 and at day 10.

### Flow cytometry analysis

To obtain single cell suspensions for flow cytometry, ventral and dorsal sheets of the ear were separated from the cartilage and incubated for 90 min at 37 °C in RPMI 1640 (Invitrogen, Grand Island, NY, USA) containing 0.01% DNAse (Sigma-Aldrich) and 0.25 mg ml^−1^ Liberase (Roche Diagnostics, Chicago, IL, USA). The digested ears were passed through a 1 ml syringe to make single-cell suspensions. The cells were filtered through 70 μm nylon mesh and washed before activation and/or staining. The following antibodies were used at 1:100 dilutions according to the manufacture’s specifications.

CD45 (30-F11, eBiosciences), CD3 (17A2, eBiosciences), CD11b (M1/70, eBiosciences), CD19 (eBioID3, eBiosciences), NK.1.1 (PK136, eBiosciences), TER-119 (Ter-119, eBiosciences), FceR1 (MAR1, eBiosciences), TCR (H57-597, eBiosciences), RoR t (B2D, eBiosciences), IL-17A (TC11-18H10, BD Pharmingen), IL-13 (eBio13A, eBiosciences), IL-5 (TFRK5, Invitrogen), and IFN-γ (XMG 1-2, eBiosciences). For intracellular cytokine staining cells were incubated for 4 hours with Leukocyte activating cocktail (BD Biosciences) in DMEM containing 2 mM L-glutamine (Invitrogen), and following surface staining cells were fixed and permeabilized according to the manufacturer’s instructions using the BD Cytofix/Cytoperm Plus Kit (BD. For counting the cells AccuCount Fluorecent particles (Spherotech, Lake Forest, IL, USA) were used. The stained cells were run on an LSR-II flow cytometer (BD Biosciences, San Jose, CA, USA) and the acquired data were analyzed using FlowJo software (Tree Star, Ashland, OR, USA).

### Statistical analysis

Mice were randomly assigned to the treatment groups and number of mice per group used in an experiment is depicted in the corresponding figure legend. Two-tailed unpaired Student’s t-test or one-way ANOVA with Tukey’s multiple comparisons were performed for significance. Mean is represented as standard error of mean (SEM) or standard deviation (SD) as shown in each figure legends. p value < 0.05 is considered as significant.

## Supplemental material

Supplemental figures can be found in the online version of this article

## Acknowledgement

We acknowledge the support of NIH (R01-AI-143790). We would like to thank Drs. C. Hunter and D. Herbert; Department of Pathology, University of Pennsylvania, PA for providing *Batf3*^*-/-*^ mice, Dr. Tiffany Scharschmidt for providing the *S. epidermidis* and Ba Nguyen for technical assistance.

## Author contributions

Conceptualization, T.P.S.; Methodology, T.P.S and P.S.; Investigation, T.P.S. and A.M.C.; Writing – original draft, T.P.S.; Writing – Review & Editing, T.P.S., E.A.C. and P.S.; Funding Acquisition, P.S. and E.A.C.; Resource, P.S. and E.A.C.; Formal analysis, T.P.S.

